# Actin limits egg aneuploidies associated with female reproductive aging

**DOI:** 10.1101/2022.05.18.491967

**Authors:** Sam Dunkley, Binyam Mogessie

## Abstract

Aging-related centromeric cohesion loss underlies premature separation of sister chromatids (PSSC) and egg aneuploidy in reproductively older females. Here we show that F-actin maintains chromatid association after cohesion deterioration in aged eggs. F-actin disruption in aged mouse eggs exacerbated PSSC, while its removal in young eggs induced extensive chromatid separation events generally only seen in advanced reproductive ages. In young eggs containing experimentally reduced cohesion, F-actin removal accelerated PSSC in a microtubule dynamics-dependent manner, suggesting that actin counteracts chromatid-pulling spindle forces. Consistently, F-actin stabilization restricted PSSC even when cohesion was acutely depleted by targeted protein degradation. We conclude that actin mitigates PSSCs arising from age-related cohesion depletion by limiting microtubule-driven chromatid separation. This is supported by a spindle-specific disruption of F-actin in aged mammalian eggs.

**One-Sentence Summary:** Actin counteracts microtubule-based pulling forces to reduce the effects of chromosome cohesion loss in aged mammalian eggs.

## Main Text

Healthy mammalian embryo formation and development critically hinges on the accuracy of sister chromatid segregation in eggs (*1, 2*). Until fertilization occurs, a tight linkage between sister chromatids’ centromeres resists their premature separation by microtubule fibers (*1, 2*). This association is predominantly achieved by Rec8-containing cohesin complexes that link sister chromatids centromeres (*1, 3–7*). Such tug-of-war between microtubule-based pulling forces and centromeric cohesion effectively generates tension between the sister chromatids’ kinetochores (*8*). Unprogrammed cohesion loss can therefore cause premature sister chromatid separation and increase the likelihood of aneuploidy (*1, 9–11*). Indeed, gradual cohesin depletion in female reproductive aging (*1, 12, 13*) is considered a major cause of egg aneuploidies broadly associated with human infertility and genetic disorders (*1, 9, 14, 15*). However, such progressive cohesion loss does not satisfactorily explain the almost exponential increase in egg aneuploidy that is observed near the end of female reproductive life (*14, 15*). Since the actin cytoskeleton is a meiosis-specific functional component of the mammalian chromosome segregation machinery (*16–19*), we addressed whether its dysfunction in eggs could explain this striking reproductive aging phenomenon.

To examine the impact of reproductive aging on cellular F-actin integrity, we first applied quantitative super-resolution microscopy and visualized fluorescent Phalloidin labelled (*17*) actin structures in eggs isolated from reproductively young (6-12 weeks old) and aged (58-62 weeks old) mice. This revealed a significant reduction in spindle F-actin fluorescence intensity in aged eggs (Fig. 1, a and b, and Fig. S1a) that did not result from decreased spindle microtubule populations (Fig. S1, b and c). This analysis further showed that cytoplasmic F-actin network density was not disrupted in aged eggs (Fig. S1, d and e). Reproductive aging in females is therefore accompanied by meiotic spindle-specific disruption of F-actin in mammalian eggs.

**Fig. 1.**
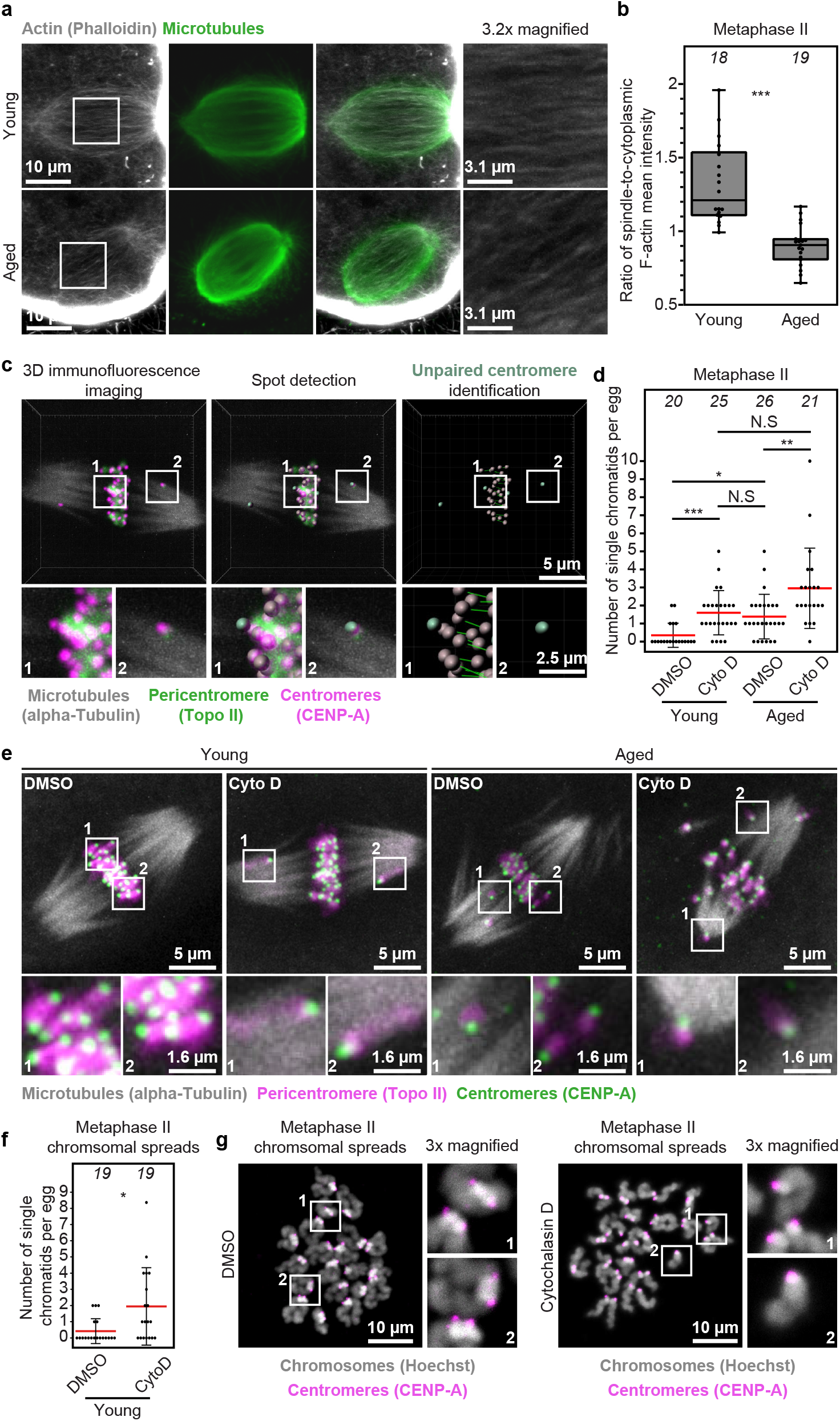
F-actin loss exacerbates reproductive age-related premature chromatid separation in mammalian eggs. (a) Sum intensity projections of phalloidin labelled spindle F-actin and microtubules in young and aged metaphase II-arrested eggs. Boxes mark regions that are magnified in insets. (b) Quantification of ratio of spindle-to-cytoplasmic F-actin mean fluorescence intensity in young and aged metaphase II-arrested eggs. Data are from 3 independent experiments. (c) Unpaired chromatid identification pipeline from maximum intensity projected immunofluorescence images of CENP-A, Topoisomerase-II and microtubules in young and aged metaphase II-arrested eggs. CENP-A spots detection and inter-centromere distance measurements in Imaris were used to identify chromatids that did not contain two centromeres and assign them as single chromatids. Boxes mark regions that are magnified in insets. (d) Quantification of the number of single chromatids (identified as in c) in DMSO- or Cytochalasin D-treated young and aged metaphase II-arrested eggs. Each filled black circle in graph represents a single egg and red bars represent mean values. Data are from 3 independent experiments. (e) Representative maximum intensity projected immunofluorescence images of microtubules, centromeres, chromatid pairs and single chromatids in DMSO- or Cytochalasin D-treated young and aged metaphase II-arrested eggs. Boxes mark regions that are magnified in insets. (f) Quantification of the number of single chromatids in metaphase II chromosomal spreads of DMSO- or Cytochalasin D-treated young eggs. Each filled black circle in graph represents chromosomal spread from a single egg and red bars represent mean values. Data are from 3 independent experiments. (g) Representative maximum intensity projected immunofluorescence images of centromeres and chromatids in metaphase II chromosomal spreads of DMSO- or Cytochalasin D-treated young eggs. Boxes mark regions that are magnified in insets. Statistical significance was evaluated using Mann-Whitney *t*-test (b and f) or Fisher’s exact test (d).

To test whether F-actin loss can exacerbate egg aneuploidies that are generally caused by gradual cohesin depletion during aging, we visualized meiotic spindles, chromosomes, and centromeres in aged eggs – isolated from reproductively aged (35-39 weeks old) mice – by high-resolution 3D immunofluorescence microscopy. We then identified sister centromere pairs using automated spot detection in Imaris (Fig. 1c) and counted prematurely separated chromatids. Consistent with previous studies of aging-related centromeric cohesion loss (*10*), this approach revealed 10/26 of DMSO-treated control aged eggs had at least two prematurely separated chromatids (Fig. 1, d and e). By stark contrast, Cytochalasin D-mediated F-actin disruption (*16, 20*) caused untimely chromatid separation in 17/21 of aged eggs (Fig. 1, d and e). These data strongly suggested that Factin limits the extent of chromatid separation in aged females, suggesting that it may mitigate the effects of age-related cohesion depletion.

To investigate the effect of F-actin removal on chromatid cohesion in young eggs, we performed similar analysis in eggs of reproductively younger (6-12 weeks old) mice. Compared to 2/20 of DMSO-treated control eggs, 12/25 of Cytochalasin D-treated young eggs displayed at least two prematurely separated chromatids (Fig. 1, d and e), which is notably comparable to incidence of chromatid separation in control aged eggs (Fig. 1e). Importantly, we were able to reproduce this high incidence of centromeric cohesion loss by treating metaphase II-arrested young eggs with Latrunculin B, a mechanistically distinct F-actin disrupting compound (*20–23*) (Fig. S2a). Similarly, quantification of chromatid linkage in independent chromosome spreads confirmed that F-actin removal predisposes eggs to centromeric cohesion loss (Fig. 1, f and g, and Fig. S2, b and c). Collectively, these results demonstrate that disruption of F-actin in young eggs is sufficient to induce a high incidence of premature sister chromatid separation similar to that seen in eggs from aged females. Notably, our quantitative immunofluorescence microscopy assays of oocyte meiosis I – when cohesion proteins can be more readily visualized along chromosome arms – did not reveal decreased Rec8-containing cohesin complexes in Cytochalasin D-treated oocytes or their corresponding chromosomal spreads (Fig. S3, a to e, and movies S1 and S2). We therefore conclude that F-actin loss leads to chromatid separation without directly disrupting canonical chromosome cohesion pathways.

To uncover how F-actin disruption accelerates chromatid separation, we implemented the TRIM-Away system (*24–31*) (Fig. 2a) to artificially deplete Rec8 in young eggs. To induce aging-like sister chromatid separation via this approach, we microinjected control IgGs or low concentrations of Rec8 heavy-chain antibodies (to only partially degrade Rec8) into TRIM21-expressing, metaphase II-arrested young eggs. High-resolution live imaging of chromatids at 5-minute intervals showed that partial Rec8 depletion in this experimental system caused modest separation and dispersion of sister chromatids (Fig. 2b, Fig. S4a, and movie S3). To quantify this, we performed 3D iso-surface reconstruction (movie S4) and object-oriented bounding box analysis in Imaris software to identify the minimal cuboid volume that contained all chromatids at each live imaging timepoint (Fig. 2b). We then plotted this volume measurement over a three-hour observation time as readout for premature separation and scattering of sister chromatids. This analysis showed that partial Rec8 degradation in DMSO-treated control eggs caused gradual sister chromatid separation with most single chromatids remaining near the spindle equator during the three-hour observation time (Fig. 2, c and d, Fig. S4b, and movie S5), and with some dispersed chromatids often re-establishing alignment with the main chromosome mass (movie S6). By contrast, partial cohesion removal after F-actin disruption dramatically accelerated chromatid separation and caused persistent scattering of single chromatids in Cytochalasin D-treated eggs (Fig. 2, c and d, Fig. S4b, and movies S7 and S8). This finding demonstrated that aging-like centromeric cohesion depletion is more likely to cause extensive aneuploidies when F-actin is absent.

**Fig. 2.**
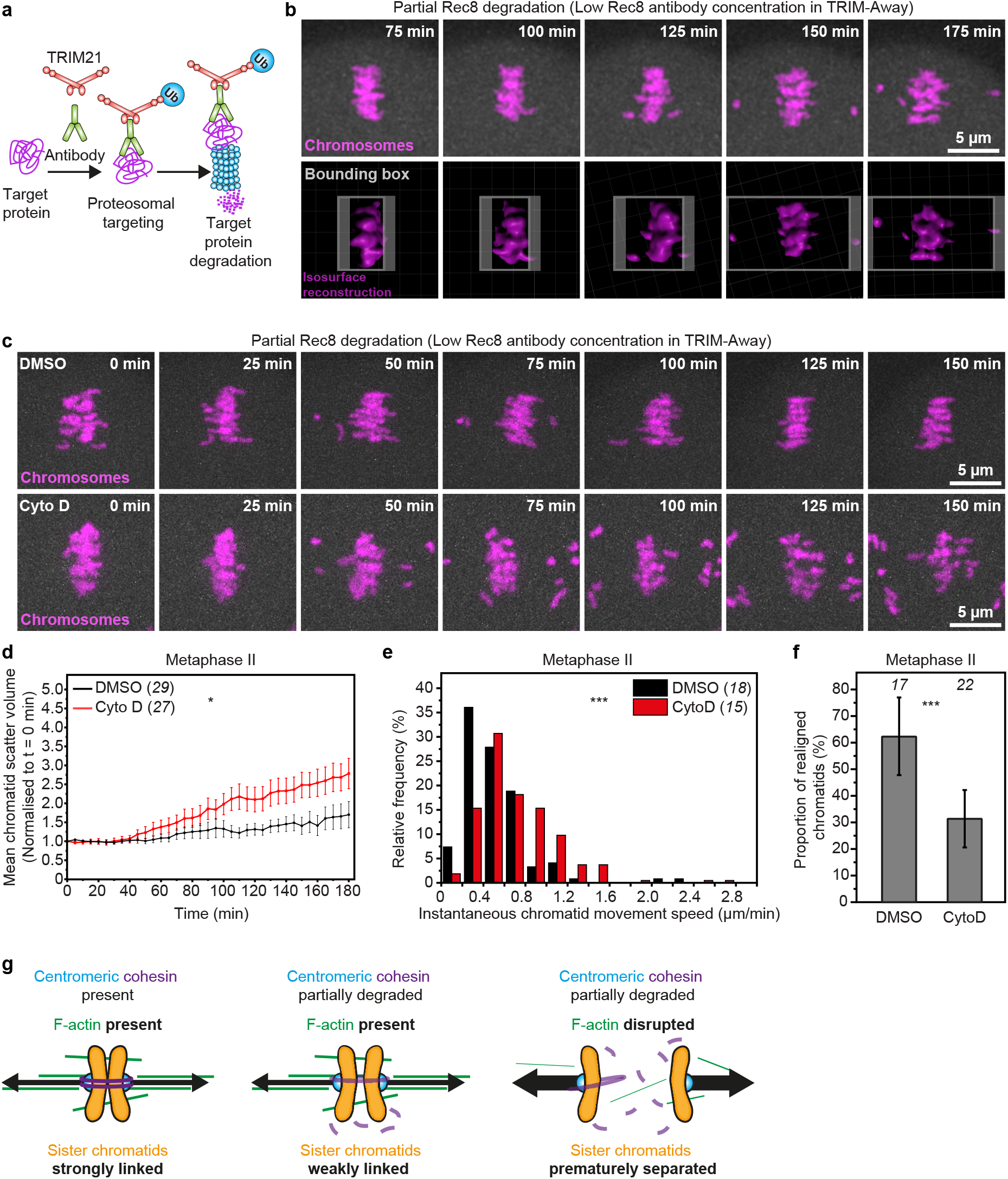
F-actin disruption in young eggs accelerates aging-like premature chromatid separation. (a) The TRIM-Away system entails microinjection of antibodies against a protein of interest and transient expression of the E3 ubiquitin ligase TRIM21, which binds Fc domain of antibodies with high affinity. TRIM21 then recruits antibody-bound target proteins for degradation by the proteasome. (b) Stills from time lapse movie of chromosomes (H2B-mRFP, top panel) and isosurface reconstructions (bottom panel) in a metaphase II-arrested egg with partially degraded Rec8. Pseudo object-orientated bounding box (grey, bottom panel) showing progressive increase in the minimal cuboid volume that encloses all chromatids as a measure of chromatid scattering. (c) Stills from representative time lapse movies of chromosomes (H2B-mRFP) in DMSO- or Cytochalasin D-treated metaphase II-arrested eggs with partially degraded Rec8. (d) Normalized chromatid scatter volumes measured as in b over 3 hours in DMSO- or Cytochalasin D-treated metaphase II-arrested eggs with partially degraded Rec8. Black and red lines represent mean values. Error bars represent S.E.M. Non-averaged individual measurements are provided in Fig. S3b. Data are from 3 independent experiments. (e) Distribution of instantaneous chromatid movement speeds in DMSO- or Cytochalasin D-treated metaphase II-arrested eggs with partially degraded Rec8. Data are from 3 independent experiments. (f) Proportion of scattered chromatids that re-established alignment to the spindle equator in DMSO- or Cytochalasin D-treated metaphase II-arrested eggs with partially degraded Rec8. Error bars represent S.D. Data are from 3 independent experiments. (g) Graphical representation of the effect of F-actin loss on chromatid linkage in eggs containing partially reduced centromeric cohesion. Statistical significance was evaluated using two-way ANOVA (d) or two-tailed Student’s *t*-test (e and f).

To test the effect of F-actin loss on chromatid motility, we tracked prematurely separated chromatids in 3D by high temporal resolution live imaging and measured their frame-to-frame displacement away from the metaphase II spindle equator. This analysis showed that most prematurely separated chromatids in Cytochalasin D-treated eggs migrated away from the spindle equator at significantly faster instantaneous speeds than in DMSO-treated control eggs that contained F-actin (Fig. 2e). This is consistent with a greater effect of chromatid-pulling microtubule forces when F-actin is disrupted. Importantly, ~62% of misaligned chromatids in control eggs became realigned at the spindle equator within the three-hours observation period (Fig. 2f). By contrast, only ~31% of previously scattered chromatids re-established correct alignment in the absence of F-actin (Fig. 2f). Persistently misaligned chromatids are more likely to be mis-segregated in anaphase II (*16*). We thus conclude that the actin cytoskeleton plays a critical, two-pronged role in the prevention of chromatid scattering: firstly, it acts as a brake against accelerated separation of sister chromatids by microtubules; secondly, it promotes the realignment of scattered single chromatids.

Having established that F-actin disruption predisposes weakly-linked sister chromatids to extensive separation (Fig. 1e), we tested whether it enrichment could help to keep them together in cohesin deficient eggs. We first stabilized F-actin in young eggs using high concentrations of SiR-Actin (*16, 17*), a fluorescence-labelled derivative of Jasplakinolide (*32*) that marks the actin cytoskeleton in living cells (*33*). We then fully degraded centromeric cohesin using highly concentrated Rec8 antibodies in our TRIM-Away strategy (Fig. 2a and movie S9). In DMSO-treated control eggs, this caused extensive separation and scattering of sister chromatids (Fig. 3, b and c, Fig. S4c, and movie S10), which is consistent with substantial depletion of cohesin from centromeric regions. In comparison, despite lack of robust cohesion, sister chromatids in SiR-Actin-treated eggs largely remained near the spindle equator during three hours of observation (Fig. 3, b and c, Fig. S4c, and movie S11). Supporting our earlier conclusion that F-actin might act as a brake against their accelerated separation, most poleward migrating chromatids also moved at significantly slower speeds when F-actin was stabilized (Fig. 3d). These data therefore collectively define a new function of F-actin in restricting the poleward movement of prematurely separated chromatids in eggs with reduced centromeric cohesion (Fig. 3e).

**Fig. 3.**
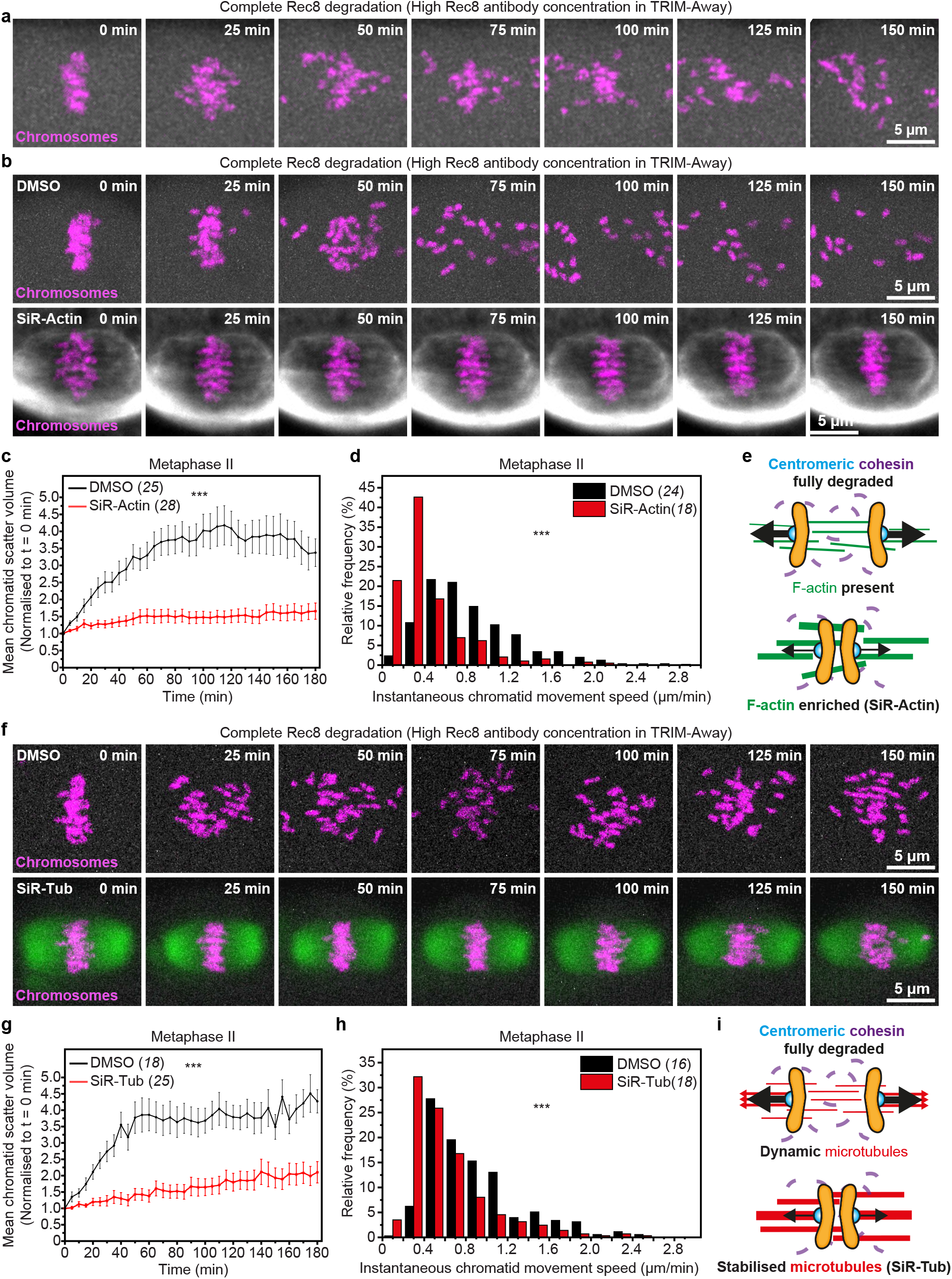
F-actin enrichment or microtubule stabilization block premature chromatid separation in the absence of centromeric cohesion. (a) Stills from time lapse movie of chromosomes (H2B-mRFP) in a metaphase II-arrested egg with fully degraded Rec8. (b) Stills from representative time lapse movies of chromosomes (H2B-mRFP) in DMSO- or SiR-Actin-treated (fluorescent F-actin shown in grey) metaphase II-arrested eggs with fully degraded Rec8. (c) Normalized chromatid scatter volumes measured as in Fig. 2b over 3 hours in DMSO- or SiR-Actin-treated metaphase II-arrested eggs with fully degraded Rec8. Black and red lines represent mean values. Error bars represent S.E.M. Non-averaged individual measurements are provided in Fig. S3c. Data are from 3 independent experiments. (d) Distribution of instantaneous chromatid movement speeds in DMSO- or SiR-Actin-treated metaphase II-arrested eggs with fully degraded Rec8. Data are from 3 independent experiments. (e) Graphical representation of the effect of F-actin stabilization on chromatid linkage in eggs with fully disrupted centromeric cohesion. (f) Stills from representative time lapse movies of chromosomes (H2B-mRFP) in DMSO- or SiR-Tubulin-treated (fluorescent microtubules shown in green) metaphase II-arrested eggs with fully degraded Rec8. (g) Normalized chromatid scatter volumes measured as in Fig. 2b over 3 hours in DMSO- or SiR-Tubulin-treated metaphase II-arrested eggs with fully degraded Rec8. Black and red lines represent mean values. Error bars represent S.E.M. Non-averaged individual measurements are provided in Fig. S3d. Data are from 3 independent experiments. (h) Distribution of instantaneous chromatid movement speeds in DMSO- or SiR-Tubulin-treated metaphase II-arrested eggs with fully degraded Rec8. Data are from 3 independent experiments. (i) Graphical representation of the effect of blocking microtubule dynamics on chromatid linkage in eggs with fully disrupted centromeric cohesion. Statistical significance was evaluated using two-way ANOVA (c and g) or two-tailed Student’s *t*-test (d and h).

Microtubules are well-established drivers of chromosomal separation during cell division (*18, 34*). Linking a decline in microtubule function with egg aneuploidy, defective microtubule dynamics were recently found to contribute to chromosomal abnormalities in aged eggs (*35*). We therefore asked whether microtubule dynamics also power chromatid movement in our experimental system of robust cohesion disruption. Consistent with earlier results, TRIM-Away-mediated acute depletion of Rec8 in DMSO-treated control eggs caused extensive dispersion of separated chromatids throughout the meiotic spindle (Fig. 3f, and movie S12). This was confirmed by bounding box analyses of chromatid mobility (Fig. 2b and 3g, and Fig. S4d). In contrast, blocking of microtubule dynamics prior to Rec8 degradation by treating eggs with high concentrations of SiR-Tubulin – a cell permeable and fluorogenic derivative of the microtubule stabilizing drug Docetaxel (*33, 36*) – significantly slowed down the movement of separated chromatids and restricted their mobility to only near the spindle equator (Fig. 3, f to h, Fig. S4d, and movie S13). These results demonstrated that microtubule dynamics drive the complete disengagement and scattering of sister chromatids following centromeric cohesion loss (Fig. 3i).

Spindle F-actin abundance directly impacts microtubule organization and dynamics in mouse eggs (*16*). To investigate whether more effective microtubule pulling in the absence of F-actin underpins chromatid separation in cohesion deficient eggs, we combined Cytochalasin D and SiR-Tubulin compounds in our experimental system of reproductive aging-like centromeric cohesion depletion (Fig. 2b). In line with our earlier data (Fig. 2, c to e), TRIM-Away-mediated partial Rec8 degradation caused rapid and extensive chromatid scattering only in eggs that did not contain F-actin (Fig. 4, a to c, Fig. S4e, and movies S14 and S15). Consistent with results showing that partial depletion of Rec8 causes only modest events of chromatid scattering (Fig. 2, c to e), SiR-Tubulin-mediated blocking of microtubule dynamics in these eggs had negligible effect on the dispersion of prematurely separated chromatids (Fig. 4, a and b, Fig. S4e, and movie S16). Similarly, the movement of unpaired chromatids was unaffected when microtubules were stabilized in eggs that were partially depleted of Rec8 (Fig. 4c). In striking contrast, blocking microtubule dynamics in eggs that did not contain F-actin – and thus experienced high incidence of chromatid scattering – significantly reduced both the extent of chromatid dispersion (Fig. 4, a and b, Fig. S4e, and movie S17) and speed of chromatid mobility (Fig. 4c). These data collectively indicate that pulling forces mediated by microtubule dynamics – normally insufficient to fully disengage weakly-bound chromatids – can cause complete chromatid separation when F-actin is lost (Fig. 4d).

**Fig. 4.**
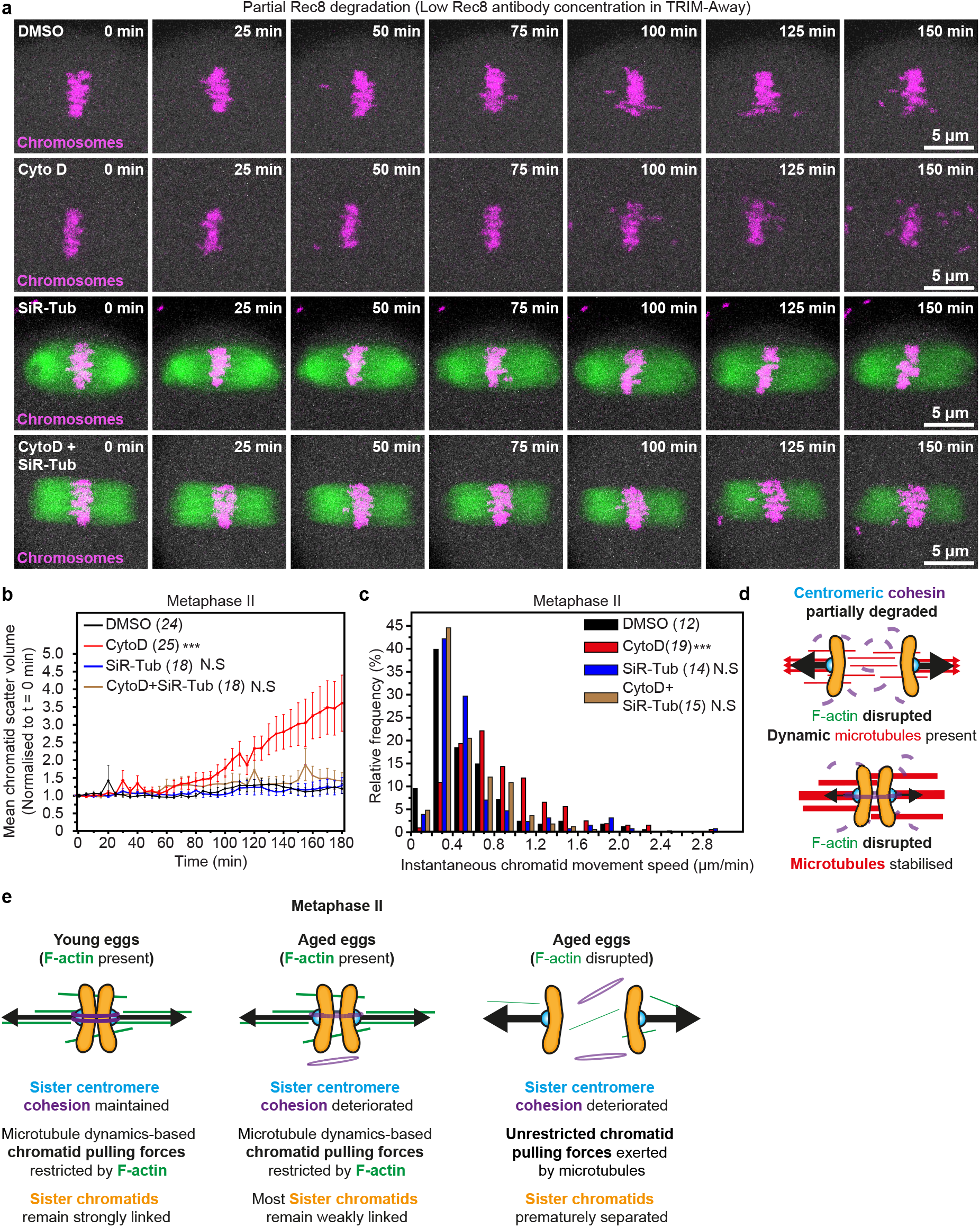
F-actin dampens microtubule-based pulling forces to prevent aging-like premature chromatid separation. (a) Stills from representative time lapse movies of chromosomes (H2B-mRFP) in DMSO-, Cytochalasin D-, SiR-Tubulin- (fluorescent microtubules shown in green) or Cytochalasin D and SiR-Tubulin-treated metaphase II-arrested eggs with partially degraded Rec8. (b) Normalized chromatid scatter volumes measured as in Fig. 2b over 3 hours in DMSO-, Cytochalasin D-, SiR-Tubulin- or Cytochalasin D and SiR-Tubulin-treated metaphase II-arrested eggs with partially degraded Rec8. Black, red, blue and brown lines represent mean values. Error bars represent S.E.M. Non-averaged individual measurements are provided in Fig. S3e. Data are from 3 independent experiments. (c) Distribution of instantaneous chromatid movement speeds in DMSO-, Cytochalasin D-, SiR-Tubulin- or Cytochalasin D and SiR-Tubulin-treated metaphase II-arrested eggs with partially degraded Rec8. Data are from 3 independent experiments. (d) Graphical representation of the effect of blocking microtubule dynamics on chromatid linkage in eggs devoid of F-actin and containing partially reduced centromeric cohesion. (e) A model for a new function of F-actin in limiting microtubule-based chromatid pulling forces in eggs of reproductively young and older females. In young eggs containing full cohesin complement, F-actin reduces the effect of chromatid pulling forces resulting in strong sister chromatid linkage. When cohesin is partially depleted in reproductive aging, F-actin-mediated resistance to microtubule pulling forces maintains linkage between most sister chromatids. When F-actin is disrupted in eggs with further reproductive aging, pulling forces generated by dynamic microtubules more effectively and prematurely separate sister chromatids. Aging-related F-actin disruption that proceeds cohesin depletion could therefore explain the dramatic rise in the incidence of egg aneuploidies near the end of female reproductive life.

Our findings indicate that spindle F-actin mitigates the effects of age-related cohesin depletion. Removal of F-actin in aged eggs with naturally reduced centromeric cohesion exacerbates premature chromatid separation. Consistently, experimentally reducing centromeric cohesion in young eggs only modestly causes untimely chromatid separation, unless F-actin is also disrupted. Because persistently misaligned chromatids in F-actin disrupted eggs are frequently missegregated in anaphase II (*16*), we propose that aging-related F-actin dysfunction can explain the sudden rise in embryo aneuploidy rate at advanced reproductive ages (*1, 14, 15*). This would be consistent with the spindle-specific physiological reduction of F-actin that we have identified in aged eggs.

Results from our combined cytoskeletal manipulation experiments indicate that F-actin exerts this previously unknown function by modulating microtubule dynamics. We previously showed that meiotic spindle F-actin stabilizes kinetochore-bound microtubules to promote accurate chromosome segregation (*16*). Furthermore, increasing the F-actin content of meiotic spindles substantially limits microtubule dynamics and reduces chromosomal separation in mouse oocytes (*16*). Our new data are consistent with a model where spindle-associated actin filaments generally dampen forces generated by dynamic microtubules (Fig. 4e). In young eggs with strongly linked chromatids, this would supplement cohesion-mediated resistance to pulling forces exerted by dynamic microtubules, thus robustly generating inter-kinetochore tension. Such force limitation would more critically resist microtubule-based chromatid pulling in aged eggs where cohesion is depleted (Fig. 4e). This would explain why F-actin disruption in aged eggs leads to extensive separation of sister chromatids. Consistent with this model, female reproductive aging is accompanied by aberrant microtubule dynamics in mouse oocytes (*35*) that are proposed as a cohesion-independent cause of meiotic aneuploidy (*37*). Thus, our findings also define F-actin disruption as a new link between cohesion deterioration and microtubule dysfunction in aging eggs.

F-actin and its associated proteins promote microtubule stability and organization in various cellular contexts, including in female meiosis (*16, 38–43*). Moreover, recent findings show that F-actin is also a component of mitotic spindle machineries that separate sister chromatids (*44, 45*). Spindle-associated actin filaments might thus have similar roles in non-reproductive cell division where aging is also associated with aneuploidies (*46–50*).

## Acknowledgments

We thank Mark Dodding for feedback on the manuscript; Kathleen Scheffler for feedback on the manuscript, cloning, advice, technical input, and TRIM-Away expertise; Michael Lampson (University of Pennsylvania) for a generous gift of Rec8 antiserum; and members of the Mogessie lab for discussions.

## Author contributions

Conceptualization: BM

Methodology: BM, SD

Investigation: SD

Visualization: BM, SD

Supervision: BM

Writing – original draft: BM

Writing – review & editing: BM, SD

## Supplementary Materials

Materials and Methods

Supplementary Text

Figs. S1 to S4

References (*51–52*)

Movies S1 to S17

## Materials and Methods

### Mouse oocyte isolation, maturation, culturing, and microinjection

All animal work in this research was performed at the University of Bristol and approved by the institution’s Animal Welfare and Ethical Review Body (AWERB). Mice were maintained in a pathogen-free environment in accordance with UK Home Office regulations under the guidelines of the University of Bristol Animal Services Unit. Oocytes were isolated from the ovaries of 8– 12-week CD1 or C57BL/6 (young) or 8-9 months CD1 or 13-14 months C57BL/6 (aged) mice, cultured, and microinjected with 6-8 picolitres of *in vitro* transcribed mRNA as described in detail recently (*17*).

C57BL/6 mice were used for comparison of cytoplasmic and spindle F-actin populations between young and aged eggs as – owing to pandemic-associated supply chain failures – this was the only aged mouse strain commercially available to us in the early stages of the project.

### Cytoskeletal drug addition experiments

Metaphase II-arrested eggs (at least 4 hours after polar body extrusion) were treated for 4 hours with: Cytochalasin D (C8273-1MG, Merck) at a final concentration of 5 μg/ml in M2 medium or Latrunculin B (428020-1MG, Merck) at a final concentration of 5 μM in M2 medium (to disrupt F-actin); SiR-Actin (SC001, Spirochrome) at a final concentration of 10 μM in M2 medium (to stabilize F-actin). In our initial experiments, we used optimized concentrations of the microtubule-stabilizing compound Taxol to effectively block microtubule dynamics in mouse eggs. However, combination of Taxol with Cytochalasin D invariably disrupted meiotic spindles. We therefore used the Docetaxel-derivative compound SiR-Tubulin (SC002, Spirochrome) at a final concentration of 1 μM in M2 medium to stabilize microtubules. In simultaneous F-actin disruption and microtubule stabilization experiments, eggs were treated with SiR-Tubulin (1 μM) for 2 hours then with a combination Cytochalasin D (5 μg/ml) and SiR-Tubulin (1 μM) for 2 hours. In TRIM-Away experiments, drug treatment was performed prior to antibody microinjection. All drugs were dissolved in DMSO (D2650-5X5ML, Merck). In control conditions, DMSO was diluted in M2 medium identically to corresponding experimental conditions.

### Generation of expression constructs and mRNA synthesis

To label chromosomes, H2B-mRFP mRNA was transcribed from pGEM-H2B-mRP(*20*). To generate pGEM-SNAP-TRIM21 for TRIM-Away experiments, pGEM-N-SNAPf was first constructed by removing SNAPf from pSNAPf (NEB, N9183S) with AgeI-XhoI and inserting it into the AgeI-XhoI site of pGEM-HE (*51*). The coding sequence of mouse TRIM21 was then transferred from pGEM-EGFP-TRIM21 (*25*) into pGEM-N-SNAPf by Gibson assembly using primers 5’TTAAACTCGAGCTCAAGCTTATGTCTCTGGAAAAGATG3’ and 5’ ATCCCGGGCCCGCGGTACCGTCACATCTTTAGTGGACAG3’.

Capped mRNAs were synthesized using a T7 polymerase (mMessage mMachine kit, Ambion). mRNA concentrations were measured using a Nanodrop spectrophotometer (Thermo Scientific).

### Fixation and immunostaining of mouse oocytes and eggs

Cells were fixed in 100 mM HEPES, 10mM MgSO_4_, 50mM EGTA, 0.5% Triton X-100 (v/v) and 2% formaldehyde (v/v) for 30 minutes at 37°C, then blocked at 4°C overnight in PBS containing 0.3% Triton X-100 (v/v) and 3% Bovine Serum Albumin (BSA) (w/v). In CENP-A immunostaining experiments, eggs were fixed at room temperature for 20 minutes, extracted in PBS containing 0.25% Triton X-100 (v/v) at room temperature for 10 minutes, and incubated in 3% BSA-PBS (w/v) at 4°C overnight. In Rec8 immunostaining staining experiments, the zona pellucida was removed before fixation by treating cells with Tyrode’s acidic solution (Merck, T1788-100ML) as described previously (*52*). In both CENP-A and Rec8 immunostaining experiments, fixed cells were incubated with λ-phosphatase (NEB, P0753S) for 2 hours at 30°C prior to immunostaining. Primary antibodies were: Rec8 rabbit antiserum (gift from Michael Lampson produced as described previously (*13*); 1:2000 dilution); CENP-A (Cell Signalling Technology, 2048S; 1:200 dilution); Topoisomerase II (Abcam, ab52934; 1:200 dilution); Human Anti-Centromere Antibody (Antibodies Incorporated, 15-234; 1:200 dilution). Secondary antibodies and stains were: Alexa-Fluor-488-labelled anti-rabbit (Molecular Probes; 1:200 dilution); Alexa-488 phalloidin (Molecular Probes; 1:20 dilution) and 5μg/ml Hoechst 33342 (Molecular Probes).

### Metaphase chromosomal spreading, fixation and immunostaining

To facilitate chromosomal spreading, the zona pellucida was removed from metaphase I oocytes or metaphase II-arrested eggs by washing cells through Tyrode’s acid solution (Merck, T1788-100ML) droplets covered with mineral oil (*52*). After washing out Tyrode’s solution in M2 medium, cells were recovered from acid treatment for a minimum of 5 minutes at 37°C. To spread chromosomes, 2-3 cells were dropped by pipetting onto a well of 15-well Multitest slide (MP Biomedicals, 096041505) containing water-based spreading solution (1% paraformaldehyde (v/v); 0.15% Triton-X100 (v/v); 3mM DTT; pH 9.2-9.4)) and air-dried in a non-transparent humidified box. After incubation with λ-phosphatase (NEB, P0753S) for 2 hours at 30°C, spreads were blocked in 3% BSA (w/v) (Fisher Scientific, 11483823) in PBS. Immunostaining was performed by washing out blocking solution in PBS followed by sequential incubation with primary and secondary antibodies for 1.5 hours at 37°C. Primary antibodies were: Rec8 rabbit antiserum (as described above and produced previously(*13*); 1:2000 dilution); Human Anti-Centromere Antibody (15-234, Antibodies Incorporated, 1:200 dilution). Secondary antibodies and stains were Alexa-Fluor-488-labelled anti-rabbit (Molecular Probes; 1:200 dilution), Alexa-Fluor-488-labelled anti-human (Molecular Probes; 1:200 dilution), and 5μg/ml Hoechst 33342 (Molecular Probes). Slides were prepared for microscopy by covering wells were Vectashield antifade mounting medium (2B Scientific, H-1000-10), mounting with 22×22 mm glass coverslips (VWR, 631-0124) and sealing with nail varnish.

### High-resolution confocal live microscopy

Confocal time lapse images of metaphase II-arrested mouse eggs were acquired using a Zeiss LSM 800 microscope equipped with an environmental chamber maintained at 37°C and a 40x C-Apochromat 1.2 NA water-immersion objective. Image acquisition using ZEN2 software (Zeiss) was performed at a temporal resolution of 5 minutes and with a Z-stack thickness of ~40 μm at 1.5 μm confocal sections. Eggs were imaged in M2 medium (with or without cytoskeletal drugs) under mineral oil as described previously (*17*).

### Confocal, super-resolution and widefield immunofluorescence microscopy

Confocal immunofluorescence images were acquired using a Zeiss LSM 800 confocal microscope, equipped with a 40x C-Apochromat 1.2 NA water immersion objective. Z-stacks were acquired with a thickness of 2.5 μm at 0.5 μm confocal sections (for spindle microtubule fluorescence quantification), 15 μm at 0.3 μm confocal sections (for single chromatid quantification) or 30 μm at 0.5 μm confocal sections (for Rec8 fluorescence quantification).

For spindle F-actin imaging, super-resolution 3D images of fluorescent phalloidin-labelled F-actin structures were acquired at the middle of the meiotic spindle in 0.5 μm steps over a range of 2.5 μm using the Airyscan module on a Zeiss LSM 800 microscope and a 40x C-Apochromat 1.2 NA water immersion objective. Post-acquisition super-resolution images were obtained by 3D Airyscan processing of raw images in ZEN2 software (Zeiss).

For cytoplasmic F-actin imaging, single section super-resolution images of fluorescent phalloidin-labelled F-actin structures were acquired at hemispheric regions of each egg using the Airyscan module on a Zeiss LSM 800 microscope and a 40x C-Apochromat 1.2 NA water immersion objective.

3D Immunofluorescence images of metaphase chromosomal spreads were acquired at 0.5 μm steps covering 5 μm using a Leica DMI6000 inverted widefield microscope equipped with a 100x HCX PL APO CS oil immersion objective.

Images in control and experimental conditions were acquired using identical imaging settings. Eggs were imaged in M2 medium under mineral oil as described previously (*17*).

### Fluorescence intensity quantification of Rec8

3D volumes of metaphase I chromosomes were reconstructed in Imaris software (Bitplane) using Hoechst 3342 immunofluorescence signal. Reconstructed surfaces were then used to mask chromosomes, remove background, and measure the mean fluorescence intensity of Rec8 on individual chromosomes.

Rec8 fluorescence intensity in metaphase I chromosomal spreads was quantified from sum intensity projections of Z-stacks (ImageJ). Mean intensity measurements were performed in ImageJ by manually drawing a region of interest around each chromosome (Fig. S3g).

Mean fluorescence intensity measurements were normalized by dividing individual values in both control and experimental groups by the average of mean fluorescence intensity values in control groups.

### Quantification of cytoplasmic and spindle F-actin fluorescence intensities in young and aged mouse eggs

The ratio of spindle F-actin to cytoplasmic F-actin fluorescence intensity was determined from sum intensity projected Z-stacks in ImageJ. Spindle F-actin fluorescence was measured by averaging the mean fluorescence intensities of 5 square regions of interest inside the spindle (Fig. S1a). Cytoplasmic F-actin fluorescence was measured by averaging the mean fluorescence intensities of 5 similarly sized square regions of interest in the area immediately surrounding the spindle (Fig. S1a). Ratios were obtained for each egg by dividing the average mean fluorescence intensity of spindle F-actin by the average mean fluorescence intensity of cytoplasmic F-actin.

Cytoplasmic F-actin fluorescence – not restricted to the region containing the meiotic spindle (Fig. S1e) – was measured in ImageJ by averaging the mean fluorescence intensities of 5 square regions of interest inside each egg. Background removal was performed by subtracting the average mean fluorescence intensity of 5 similarly sized square regions of interest – placed outside the image of the cell and in regions that did not contain any phalloidin signal – from the averaged mean fluorescence intensity of cytoplasmic F-actin (Fig. S1e). Mean fluorescence intensity measurements were normalized by dividing individual values in both control and experimental groups by the average of mean fluorescence intensity values in control groups.

### Quantification of spindle microtubule fluorescence intensity in young and aged mouse eggs

Spindle microtubule fluorescence was measured in ImageJ from sum intensity projected Z-stacks by averaging the mean fluorescence intensities of 5 square regions of interest inside the spindle (Fig. S1b). Background removal was performed by subtracting the average mean fluorescence intensity of 5 similarly sized square regions of interest – placed in cytoplasmic regions that did not contain any microtubule filament – from the averaged mean fluorescence intensity of spindle microtubules (Fig. S1b). Mean fluorescence intensity measurements were normalized by dividing individual values in both control and experimental groups by the average of mean fluorescence intensity values in control groups.

### Identification and quantification of prematurely separated sister chromatids

Chromatids in metaphase II-arrested eggs were identified by reconstructing the 3D volume of Topoisomerase II immunofluorescence signal in Imaris software (Bitplane). Centromeres were then identified using CENP-A immunofluorescence signal for spot detection in Imaris. Lastly, neighboring centromeres were manually assigned as sisters using distance measurements and reconstructed chromatid volumes as reference. Chromatids containing centromeres for which no corresponding sisters were identified via this analysis were designated as prematurely separated chromatids (Fig. 1c). Single chromatids in metaphase II spreads were identified via Hoechst labelling and centromere distance measurements in combination with spot detection in Imaris using the immunofluorescence signal of human anti-centromere antibody.

### Partial and complete targeted degradation of Rec8 in TRIM-Away experiments

In partial Rec8 degradation TRIM-Away experiments, TRIM21 expressing, metaphase II-arrested eggs were microinjected with 2-3 picolitres of Rec8 antiserum (*13*) (1:30-1:50 dilution) and Alexa Fluor 488 Dextran 10,000 MW (to validate antibody microinjection by fluorescence imaging) (Molecular Probes, D22910; 1:40 dilution) in 0.05% (v/v) NP40-PBS. In complete Rec8 degradation TRIM-Away experiments, TRIM21 expressing, metaphase II-arrested eggs were microinjected with 2-3 picolitres of Rec8 antiserum (*13*) (1:2 dilution) and Alexa Fluor 488 Dextran 10,000 MW (Molecular Probes, D22910; 1:40 dilution) in 0.05% (v/v) NP40-PBS. In simultaneous Rec8 degradation and cytoskeletal manipulation experiments, all TRIM-Away microinjections were performed in M2 medium containing individual or a combination of cytoskeletal drugs as appropriate.

### 3D chromatid surface reconstruction and scattering volume measurement

Chromatid surfaces were reconstructed in 3D in Imaris (Bitplane) from high-resolution time lapse movies of H2B-mRFP. Object-oriented bounding box analysis was performed in Imaris to identify the minimal cuboid volume that enclosed all chromatids at each time point. This bounding box volume was defined as the chromatid scattering volume. For each egg, data normalization was performed by dividing measured values at each time point with the bounding box volume at the start of the live imaging experiment (t = 0 mins).

### Quantification of 3D chromatid movement speed and realignment

To quantify chromatid mobility, the 3D volumes of chromatids were reconstructed in Imaris (Bitplane) from high-resolution live imaging datasets of H2B-mRFP. For chromatids that distinctly migrated away from the spindle equator in control and experimental conditions, frame-to-frame displacement was measured by tracking the position of chromatid leading ends via the Measurement Points function in Imaris. Displacement values (μm) were divided by the temporal resolution of live imaging experiments (5 minutes) to calculate instantaneous chromatid movement speeds.

Realignment of previously scattered single chromatids to the spindle equator was analyzed in 3D in Imaris. A realignment event was defined as the merging of a misaligned chromatid with the main chromosome mass for a duration of at least 10 minutes.

### Statistical data analyses

Histograms, box plots and other graphs were generated using OriginPro (OriginLab) or Prism (GraphPad) software. Box plots show mean (line), 5^th^, 95^th^ (Whiskers), 25^th^ and 75^th^ percentile (box enclosing 50% of the data) and are overlaid with individual data points. Statistical significance evaluations, one-way and two-way analyses of variance were performed in OriginPro or Prism software. Significance values are \**P*<0.05, \*\**P*<0.005 and \*\*\**P*<0.0005. Nonsignificant values are indicated as N.S.

---

**Fig. S1.**
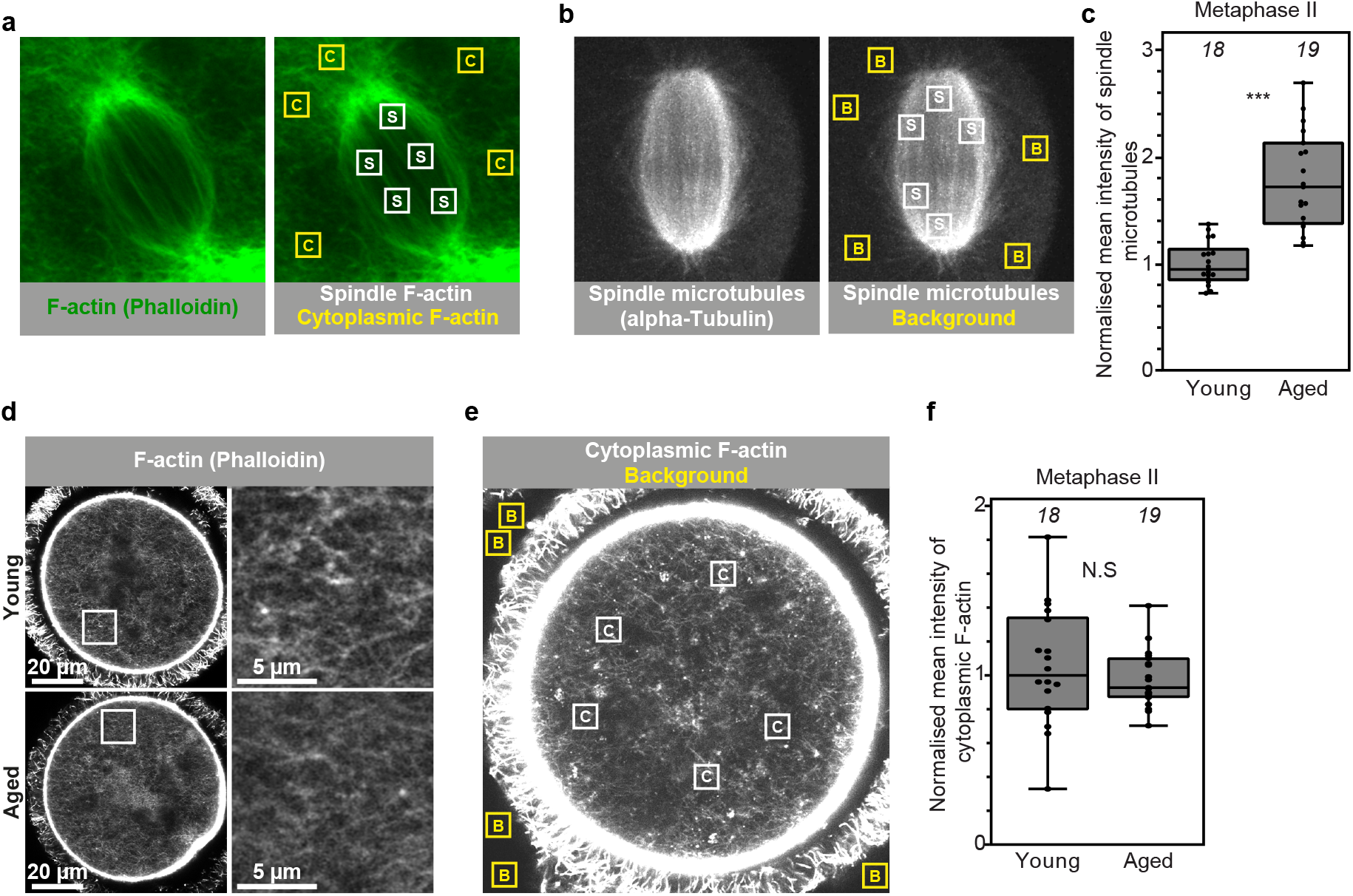
Female reproductive aging is accompanied by spindle-specific F-actin loss in eggs. (a) Method (described in Materials and Methods) for quantification of spindle-to-cytoplasmic F-actin mean fluorescence intensity ratio in metaphase II-arrested eggs of reproductively young or aged mice. (b) Method (described in Materials and Methods) for quantification, background correction and normalization of spindle microtubule mean fluorescence intensities in eggs of reproductively young or aged mice. (c) Normalized spindle microtubule mean fluorescence intensities in young and aged metaphase II-arrested eggs. Data are from 3 independent experiments. (d) Representative single section Airyscan images of phalloidin labelled cytoplasmic F-actin structures in metaphase II-arrested eggs of reproductively young or aged mice. Boxes mark regions that are magnified in insets. (e) Method (described in Materials and Methods) for quantification, background correction and normalization of cytoplasmic F-actin mean fluorescence intensities in eggs of reproductively young or aged mice. (f) Normalized cytoplasmic F-actin mean fluorescence intensities in young and aged metaphase II-arrested eggs. Data are from 3 independent experiments. Statistical significance was evaluated using Mann-Whitney *t*-test (c and f).

**Fig. S2.**
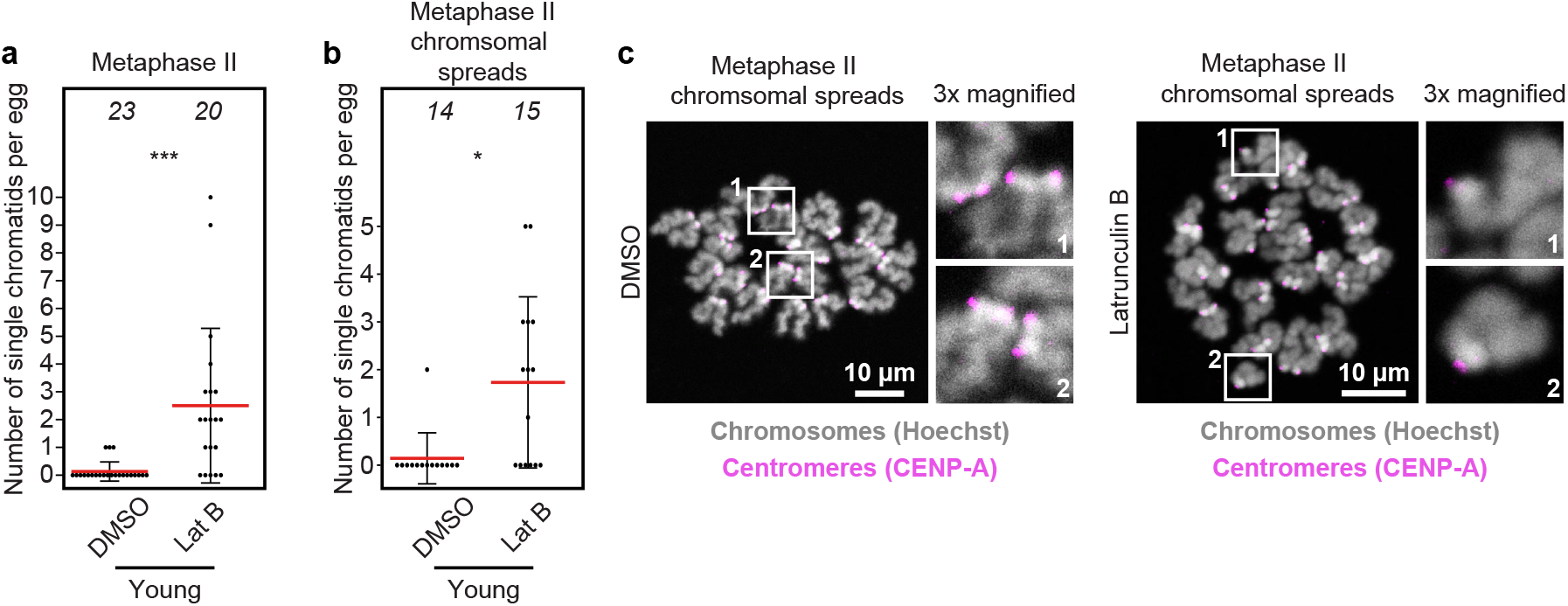
F-actin disruption predisposes young eggs to premature sister chromatid separation. (a) Quantification of the number of single chromatids (identified as in Fig. 1c) in DMSO- or Latrunculin B-treated metaphase II-arrested eggs. Each filled black circle in graph represents a single egg and red bars represent mean values. Data are from 3 independent experiments. (b) Quantification of the number of single chromatids in metaphase II chromosomal spreads of DMSO- or Latrunculin B-treated eggs. Each filled black circle in graph represents chromosomal spread from a single egg and red bars represent mean values. Data are from 3 independent experiments. (c) Representative maximum intensity projected immunofluorescence images of centromeres and chromatids in metaphase II chromosomal spreads of DMSO- or Latrunculin B-treated young eggs. Boxes mark regions that are magnified in insets. Statistical significance was evaluated using Mann-Whitney *t*-test (a and b).

**Fig. S3.**
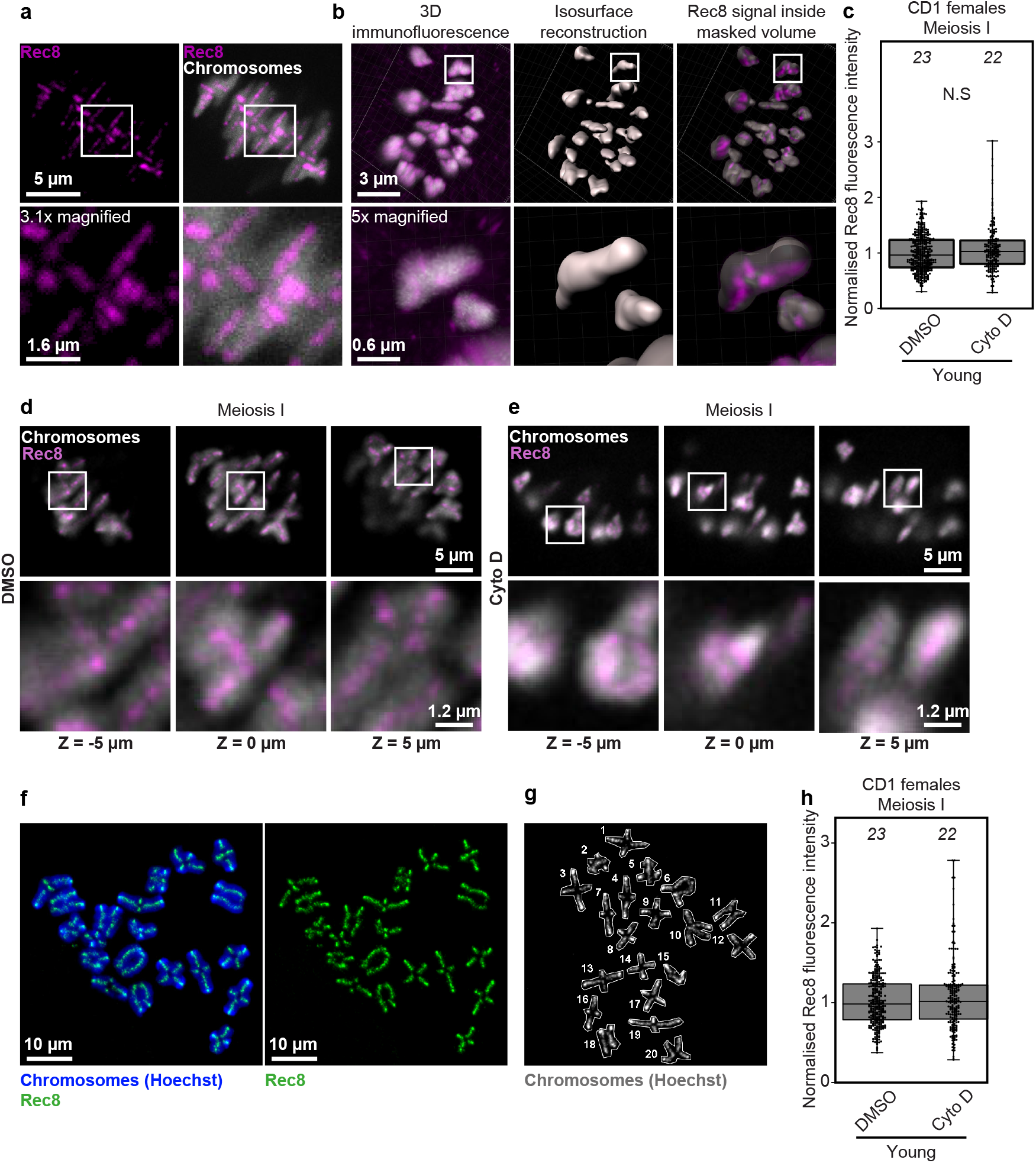
F-actin disruption does not impact classical mechanisms of centromeric cohesion. (a) Maximum intensity projected high-resolution immunofluorescence images of Rec8 cohesin complexes and homologous chromosomes in a mouse oocyte. Boxes mark regions that are magnified in insets. (b) Rec8 immunofluorescence intensity quantification pipeline. Individual chromosome volumes were reconstructed using the Surfaces module of Imaris. Mean Rec8 fluorescence intensity was then measured inside the masked volume each chromosome. (c) Normalized Rec8 mean fluorescence intensities in DMSO- or Cytochalasin D-treated mouse oocytes. Data are from 3 independent experiments. (d) Representative single confocal section immunofluorescence images of Rec8 and chromosomes – spaced 5 μm apart – in DMSO-treated mouse oocytes. Boxes mark regions that are magnified in insets. (e) Representative single confocal section immunofluorescence images of Rec8 and chromosomes – spaced 5 μm apart – in Cytochalasin D-treated mouse oocytes. Boxes mark regions that are magnified in insets. (f) Representative maximum intensity projected immunofluorescence images of Rec8 and chromosomes in metaphase I chromosomal spread of a mouse oocyte. (g) Method (described in Materials and Methods) for quantification of Rec8 mean fluorescence intensity in metaphase I chromosomal spreads of mouse oocytes. (h) Normalized Rec8 mean fluorescence intensities in metaphase I chromosomal spreads of DMSO- or Cytochalasin D-treated mouse oocytes. Data are from 3 independent experiments. Statistical significance was evaluated using Mann-Whitney (c) and Welch’s (h) *t*-test.

**Fig. S4.**
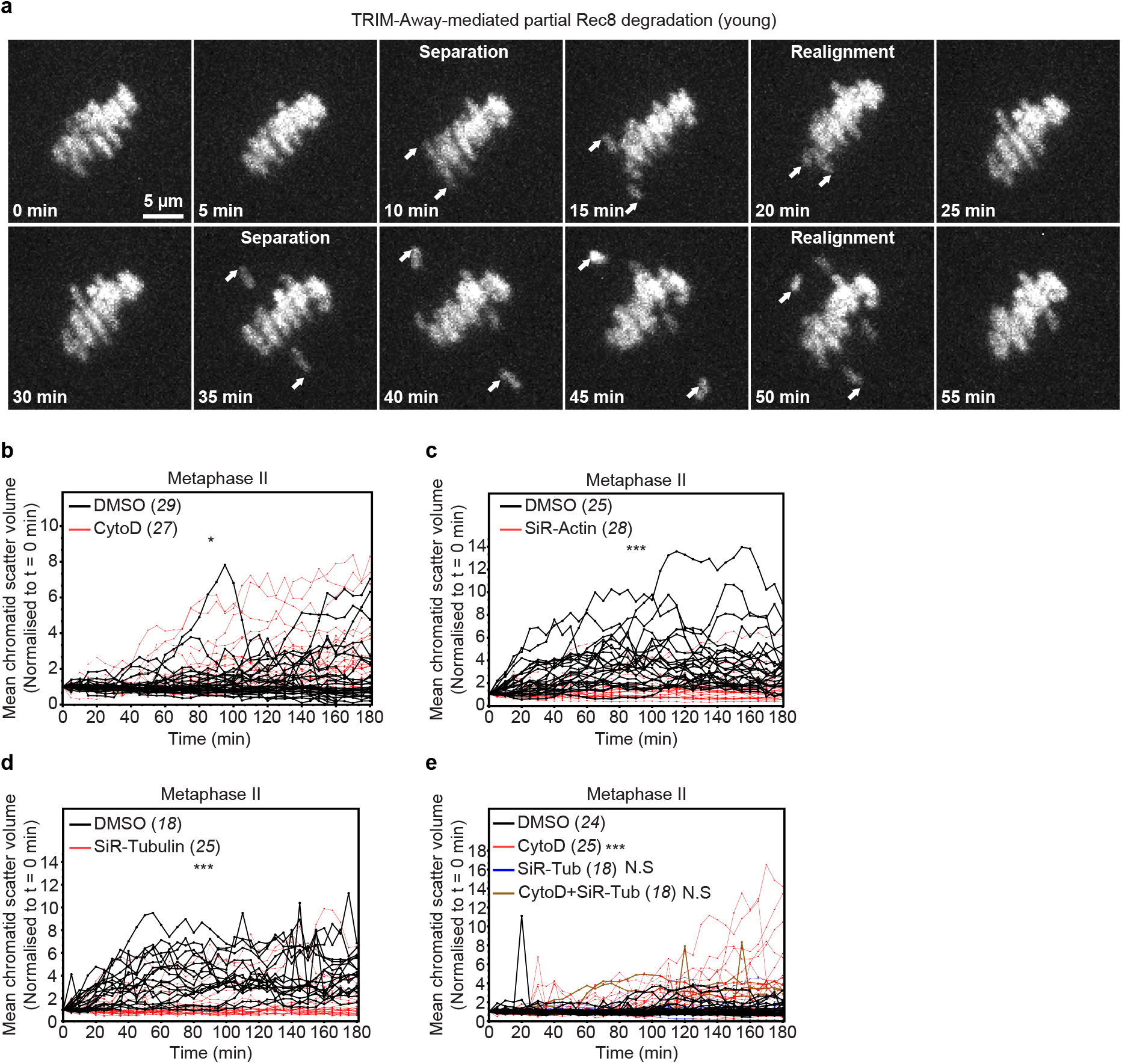
F-actin limits aging-like premature chromatid separation by microtubule-based pulling forces. (a) Representative maximum intensity projected high-resolution confocal images of sister chromatids in a metaphase II-arrested mouse egg with partially degraded Rec8. Arrows indicate modest chromatid separation and subsequent realignment events. (b) Normalized chromatid scatter volumes measured as in Fig. 2b over 3 hours in DMSO- or Cytochalasin D-treated metaphase II-arrested eggs with partially degraded Rec8. Each black and red line represents measurements from a single egg. Averaged measurements are presented in Fig. 2d. Data are from 3 independent experiments. (c) Normalized chromatid scatter volumes measured as in Fig. 2b over 3 hours in DMSO- or SiR-Actin-treated metaphase II-arrested eggs with fully degraded Rec8. Each black and red line represents measurements from a single egg. Averaged measurements are presented in Fig. 3c. Data are from 3 independent experiments. (d) Normalized chromatid scatter volumes measured as in Fig. 2b over 3 hours in DMSO- or SiR-Tubulin-treated metaphase II-arrested eggs with fully degraded Rec8. Each black and red line represents measurements from a single egg. Averaged measurements are presented in Fig. 3g. Data are from 3 independent experiments. (e) Normalized chromatid scatter volumes measured as in Fig. 2b over 3 hours in DMSO-, Cytochalasin D-, SiR-Tubulin- or Cytochalasin D and SiR-Tubulin-treated metaphase II-arrested eggs with partially degraded Rec8. Each black and red line represents measurements from a single egg. Averaged measurements are presented in Fig. 4b. Data are from 3 independent experiments.

**Movie S1.** Navigation through single immunofluorescence confocal sections – spaced 0.5 μm apart – of Rec8 (magenta) and homologous chromosomes (grey) in a DMSO-treated mouse oocyte.

**Movie S2.** Navigation through single immunofluorescence confocal sections – spaced 0.5 μm apart – of Rec8 (magenta) and homologous chromosomes (grey) in a Cytochalasin D-treated mouse oocyte.

**Movie S3.** Time lapse movie of modest separation of chromatids (H2B-mRFP, magenta) in a metaphase II-arrested mouse egg with partially degraded Rec8.

**Movie S4.** 3D isosurface reconstruction (Imaris) of chromatids in movie 3 for bounding box analysis of chromatid scattering.

**Movie S5.** Time lapse movie of modest separation of chromatids (H2B-mRFP, magenta) in a DMSO-treated metaphase II-arrested mouse egg with partially degraded Rec8.

**Movie S6.** Time lapse movie of modest separation and subsequent realignment (indicated by write arrows) of chromatids (H2B-mRFP, magenta) in a DMSO-treated metaphase II-arrested mouse egg with partially degraded Rec8.

**Movie S7.** Time lapse movie of excessive separation and scattering of chromatids (H2B-mRFP, magenta) in a Cytochalasin D-treated metaphase II-arrested mouse egg with partially degraded Rec8 (example 1).

**Movie S8.** Time lapse movie of excessive separation and scattering of chromatids (H2B-mRFP, magenta) in a Cytochalasin D-treated metaphase II-arrested mouse egg with partially degraded Rec8 (example 2).

**Movie S9.** Time lapse movie of complete separation and scattering of chromatids (H2B-mRFP, magenta) in a metaphase II-arrested mouse egg with fully degraded Rec8.

**Movie S10.** Time lapse movie of complete separation and scattering of chromatids (H2B-mRFP, magenta) in a DMSO-treated (control for SiR-Actin treatment) metaphase II-arrested mouse egg with fully degraded Rec8.

**Movie S11.** Time lapse movie of restricted movement of chromatids (H2B-mRFP, magenta) in a SiR-Actin-treated metaphase II-arrested mouse egg with fully degraded Rec8. SiR-Actin fluorescence in shown in grey.

**Movie S12.** Time lapse movie of complete separation and scattering of chromatids (H2B-mRFP, magenta) in a DMSO-treated (control for SiR-Tubulin treatment) metaphase II-arrested mouse egg with fully degraded Rec8.

**Movie S13.** Time lapse movie of restricted movement of chromatids (H2B-mRFP, magenta) in a SiR-Tubulin-treated metaphase II-arrested mouse egg with fully degraded Rec8. SiR-Tubulin fluorescence in shown in green.

**Movie S14.** Time lapse movie of complete separation and scattering of chromatids (H2B-mRFP, magenta) in a DMSO-treated (control for Cytochalasin D + SiR-Tubulin treatment) metaphase II-arrested mouse egg with partially degraded Rec8.

**Movie S15.** Time lapse movie of excessive separation and scattering of chromatids (H2B-mRFP, magenta) in a Cytochalasin D-treated (control for Cytochalasin D + SiR-Tubulin treatment) metaphase II-arrested mouse egg with partially degraded Rec8.

**Movie S16.** Time lapse movie of restricted movement of chromatids (H2B-mRFP, magenta) in a SiR-Tubulin-treated (control for Cytochalasin D + SiR-Tubulin treatment) metaphase II-arrested mouse egg with partially degraded Rec8. SiR-Tubulin fluorescence in shown in green.

**Movie S17.** Time lapse movie of restricted movement of chromatids (H2B-mRFP, magenta) in a Cytochalasin D + SiR-Tubulin-treated metaphase II-arrested mouse egg with partially degraded Rec8. SiR-Tubulin fluorescence in shown in green.

